# Integrating the host plant phylogeny, plant traits, intraspecific competition and repeated measures using a phylogenetic mixed model of field behaviour by polyphagous herbivores, the leaf-cutting ants

**DOI:** 10.1101/854471

**Authors:** Manasee Weerathunga, Alexander S. Mikheyev

**Affiliations:** Ecology and Evolution Unit, Okinawa Institute of Science and Technology Graduate University, Onna-son, Tancha, Okinawa, Japan; Evolutionary Genomics Research Group, Australian National University, Canberra, Australian Capital Territory, Australia

**Keywords:** foraging, leaf-cutting ants, neotropics, palatability, phylogenetic models, PGLS

## Abstract

Herbivores use a wide range of factors to choose their host, including their own physiological states, physical characteristics of plants, and the degree of competition. Field observations of herbivores in their native habitats provide a means for simultaneously estimating the relative importance of these factors, but statistical analysis of all these factors may be challenging. Here we used a 7-wk dataset of leaf-cutting ant (*Atta cephalotes*) foraging in a diverse neotropical arboretum containing 193 tree species (822 trees) to examine the relative role of tree phylogeny, territoriality and tree functional characteristics using a phylogenetic generalized least squares (PGLS) model. We observed that 54 tree species (117 trees) were foraged by the ants. This pattern was not random, but reflected known features of leaf-cutting ant foraging behaviour, such as the preference for larger trees and the decreased likelihood of foraging at the periphery of a colony’s territory. However, random effects such as tree phylogeny, the identity of individual trees and colony-specific effects, explained most of the variation in foraging data. A significant phylogenetic effect on foraging likelihood (λ = 0.28), together with repeated measures of foraging on the same tree species, allowed estimation of relative palatability for each plant species. PGLS models can be flexibly scaled to include other covariates for even more complex investigation of foraging behaviour, and the link function can be modified to include the amount of plants foraged. As a result, PGLS can be used as a flexible framework for the study of LCA foraging.

## Introduction

Plant-herbivore interactions play a major role in structuring ecosystems by influencing competitive ability of plants, cycling biomass in ecosystems, and energy transfer to higher trophic levels (Hunter & Watt 2008). Herbivores are considered keystone species, where attributes such as consumption intensity of herbivores, herbivore abundance and the herbivore diversity act as determinants of meta-community structure (Carson & Root 2000, Smith *et al*. 2010, Wan *et al*. 2015). In addition to altering plant diversity and abundance, herbivory also drives the evolution of plant defences (Poelman & Kessler 2016). On the other hand, phylogenetic composition of host plant communities influences evolution of herbivore specialization, and herbivore biodiversity depends on their coevolutionary interactions with plants (Volf *et al*. 2017). As a result, herbivores show increased preference towards congeneric and confamilial hosts (Janz & Nylin 1998, Novotny *et al*. 2002, Winkler & Mitter 2008), making host plant phylogeny an important clue in predicting herbivore foraging preference, food webs and community structure (Volf *et al*. 2015, 2017, Winkler & Mitter 2008).

While many herbivores are highly host-specific, others consume a wide variety of plant taxa. Much work has gone into understanding how polyphagous herbivores make their choices, particularly using field observations. However, statistical analysis of field observations is not straightforward for a number of reasons. First, host plants are not phylogenetically independent, requiring some sort of approach to control for phylogeny. Second, the analyses often contain both fixed and random effects in analyses. Fixed effects refer to terms in which the group mean is fixed at a set number of levels, they often correspond to specific experimental conditions or ecological categories (e.g. leaf toughness and habitat type). By contrast, random effect means are assumed to be drawn from a large underlying populations. They include effects of individuals measured repeatedly and multiple randomly chosen sites. Phylogenetic generalized least squares (PGLS) models can incorporate phylogenies and deal with mixed (fixed and random) effects providing a solution to this problem (Bürkner 2017, Covarrubias-Pazaran 2016, Hadfield 2010). Here, we apply PGLS to a well-studied group dominant polyphagous tropical herbivores, the leaf-cutting ants.

Leaf cutting ants (LCA) are one of the most dominant herbivores in thew neotropics (Hölldobler & Wilson 1990, Meyer *et al*. 2011) capable of processing a substantial proportion (50%->90%) of plant species within their habitat (Rockwood 1976, Shepherd 1985, Vasconcelos 1997, Wirth *et al*. 1997). While LCA are polyphagous they are also selective in the kinds of plants they consume and how much material they take. Their herbivory has measurable ecosystem-wide effects making them one of the most prominent ecosystem engineers in the neotropics (Costa *et al*. 2008) and potentially dangerous invasive species (Mikheyev 2007). The selectivity shown by LCA towards various plant species plays a significant role in this. Plant species preferred by LCA are likely to decline at a rapid rate while other species would proliferate, thus altering the community structure (Farji-Brener *et al*. 2000). For example, in savanna ecosystems LCA partially defoliation less-preferred species and completely defoliate of highly-preferred species (Costa *et al*. 2018). LCA also account for a prominent negative effect on forest regrowth by harvesting 12%-17% of annual forest reproduction (Cherrett 1986). Similarly, in savanna ecosystems they are known to act as an ecological filter affecting the structure and composition of vegetation, creating demographic bottlenecks by reducing seed availability and seedling survival (Costa *et al*. 2017). In both these cases, the plant selectivity shown by LCA will play a prominent role ecosystem engineering pattern, since the preferred species will be removed from the community at a higher rate. Hence, LCA prove to be an ideal model to study the top-down pressure exerted by herbivores on engineering Neotropical community structure and functions, and their foraging preferences have been extensively studied. As a result we can ask whether the proposed statistical framework recovers some key attributes already known about LCA foraging.

Research shows that foraging ants always have to balance the energetic costs involved in foraging with the quality, quantity and suitability of harvested material (Olsson *et al*.2008) while balancing the fulfillment of the nutrient requirement and minimizing toxic intake (Mundim *et al*. 2009). Foraging behavior of LCA commonly involve foraging in palatable patches, especially in patches in close vicinity of colonies. Such preferred sites are known to depend on the preferences of founding queens (Vasconcelos & De Vasconcelos 1990). This possibly plays an important role in ecosystem engineering, because extensive foraging in palatable patches results in gaps in the canopy and changes in plant performance and viability in those patches, leading to changes in forest structure and microclimate (Meyer *et al*. 2011). Thus, the question arises whether extensive herbivory on preferred patches based on foraging preferences play a role in alteration of ecosystem structure, plant performance, microclimate and resource availability (Meyer *et al*. 2011).

The foraging strategies of LCA differ broadly with the quality, quantity, suitability and availability of plant substrate, leading diet breadth to vary widely with spatio-temporal availability (Costa *et al*. 2018). Sometimes the foraging preference of LCA is determined by synergistic effect of two or more factors mentioned above or by the isolated impact of one of those (Berish 1986, Bowers & Porter 1981, Howard 1987). However, despite the extensive research on LCA foraging preferences, it has been hard to integrate a single model that would test the synergistic effect of all possible factors owing to the paucity in research simultaneously discussing the influence of all possible factors. In a recent advance, Gerhold *et al*. (2019) integrated the phylogenetic signal in LCA diet, which allowed them to control for plant phylogeny in the analysis of leaf toughness. However, their model did not allow them to incorporate other factors influencing ant foraging, such as colony-level effects and effect of territoriality (competition) on LCA diet.

In this study we generalized these findings by using a phylogenetically controlled mixed model approach to simultaneously determine the impact of fixed effects such as resource availability, distance from the nest and effects such as phylogenetic signal and colony-level effects on LCA diet. We found that this approach captures known patterns of LCA foraging, but also indicates the surprisingly large role of intraspecific effects.

## Material and Methods

### Field surveys

Our study took place in the Holdridge Arboretum of La Selva Biological Station, Heredia Province, Costa Rica. Management consisted of regular mowing of grass and shrub removal, so that most of the biomass harvestable by the ants was at canopy level. We recorded the foraging patterns of eight mature *A. cephalotes* individuals at night during times of peak leaf-cutting activity on a nightly basis over the course of 7 wk during the dry season from mid-February to late April 2003. The observations consisted of systematically following every active trail of *A. cephalotes* foragers from the mound to the trees on which they were foraging. We identified the trees by recording tree tags or, when tags were missing, by inferring their identity using the arboretum database from size and the relative position of nearby trees. An *A. cephalotes* territory was defined as the minimum convex polygon containing all the trees harvested by the colony, which were calculated using the Animal Movement extension for Arcview. In the course of the study, we observed the ants cutting 63 out of the 225 species present in the arboretum (28%) (Figure 1).

**Figure 1.**
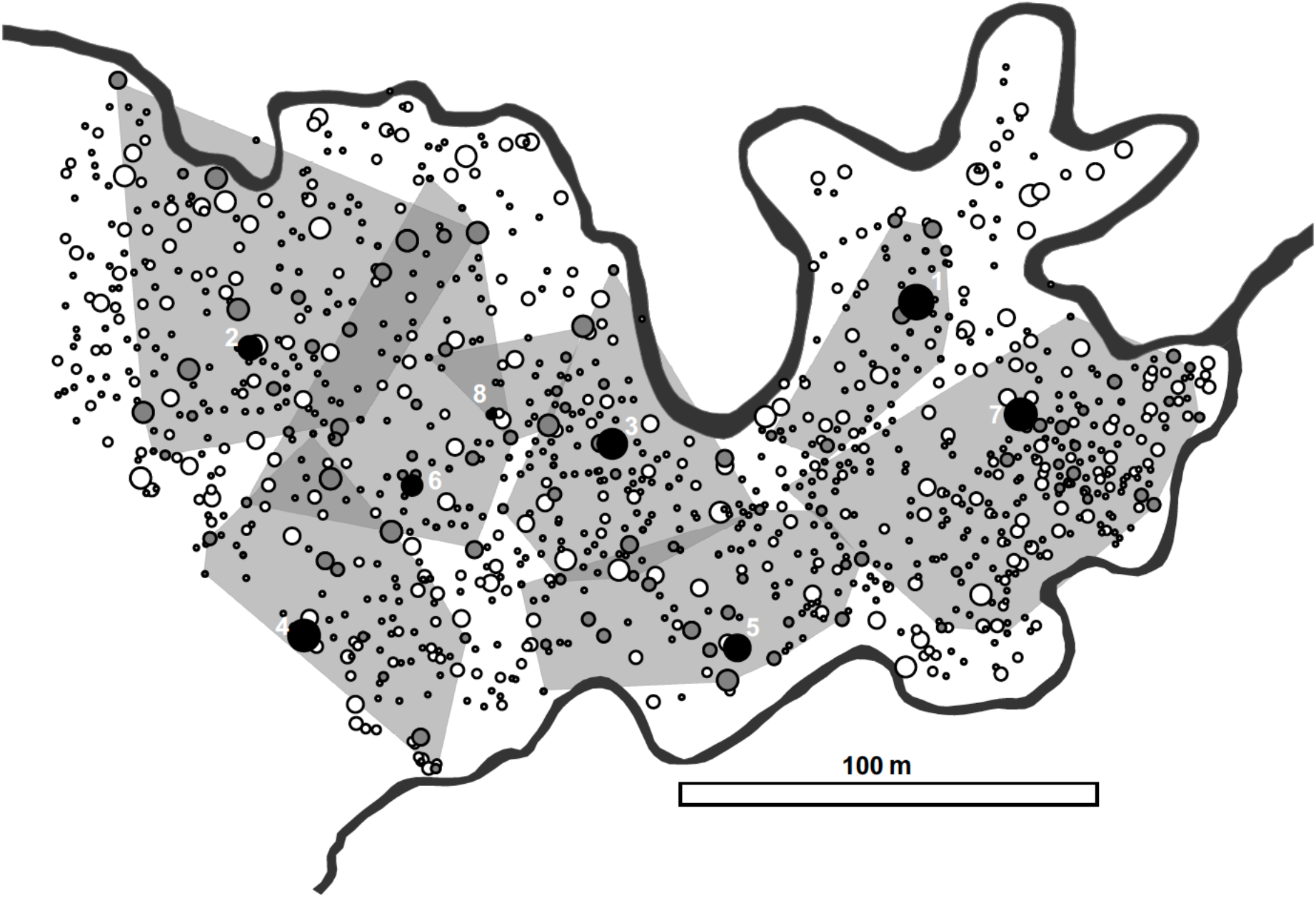
Foraging territories of *Atta cephalotes* colonies in the Holdridge arboretum (La Selva, Costa Rica). Actual sizes or mounds are shown by black circles and numbered. The river bounding the arboretum is shown as a black line. Trees are shown as either open (non-foraged) or grey (foraged) circes. Territories are shown as grey polygons. The territory boundaries shifted in the course of the experiment, often accompanied by battles between the colonies. As a result, some of the trees were in overlapping territories and could have been consumed by multiple colonies over the course of the study. While the Arboretum is unusually species-rich, even for a tropical forest, many species were present multiple times and repeated measures of palatability were possible.

### Statistical analysis

Our overall goal was to implement a statistical model that could flexibly account for fixed and random effects affecting observed LCA foraging patterns. We used a Bayesian logistic mixed model implemented *brms* package (Bürkner 2017), which can incorporate a range of covariates, including the phylogeny of the plant species. This statistical framework can also incorporate included ‘fixed’ effects, namely tree size (diameter at breast height (dbh), and tree distance from the focal colony. These effects contain a defined number of levels within the experiment. In this case, the fixed effects corresponded to known features of leaf cutting ant foraging, which we hoped to recover using the model as a way of validating this approach. However, like many other PGLS software packages, *brms* can also handle ‘random’ effects, namely the identity of an individual tree, as well as colony-level and phylogenetic effects.

Random effects have many levels, and not all of them are captured in the experimental design. In our design, factors like the identity of individual trees could encompass numerous unmeasured factors, such as its phenology, age, physiological state, *etc*. However, because all of the effects are present in the same model, we can compare the relative contribution of fixed vs random effects, and to examine the effects separately within the same data set to compare their relative explanatory power. In principle, any number of additional factors could be added to the model, given enough available data.

For the phylogenetic covariate we used Phylomatic (Webb & Donoghue 2005) to extract phylogenetic data for as many tree species as possible. We were able to place 193 out of 225 species onto the Zanne *et al*. (2014) reference and include them for further analysis. This tree was used to compute a variance-covariance matrix that was added as a random effect to the model.

### Data accessibility

The full details of the statistical analysis, the raw data and all steps necessary to reproduce summary statistics and the main data plots are available in the git repository https://github.com/MikheyevLab/LCA-foraging. A web page with all of the analyses and code used to generate Figure 2 can be viewed at https://mikheyevlab.github.io/LCA-foraging/.

**Figure 2.**
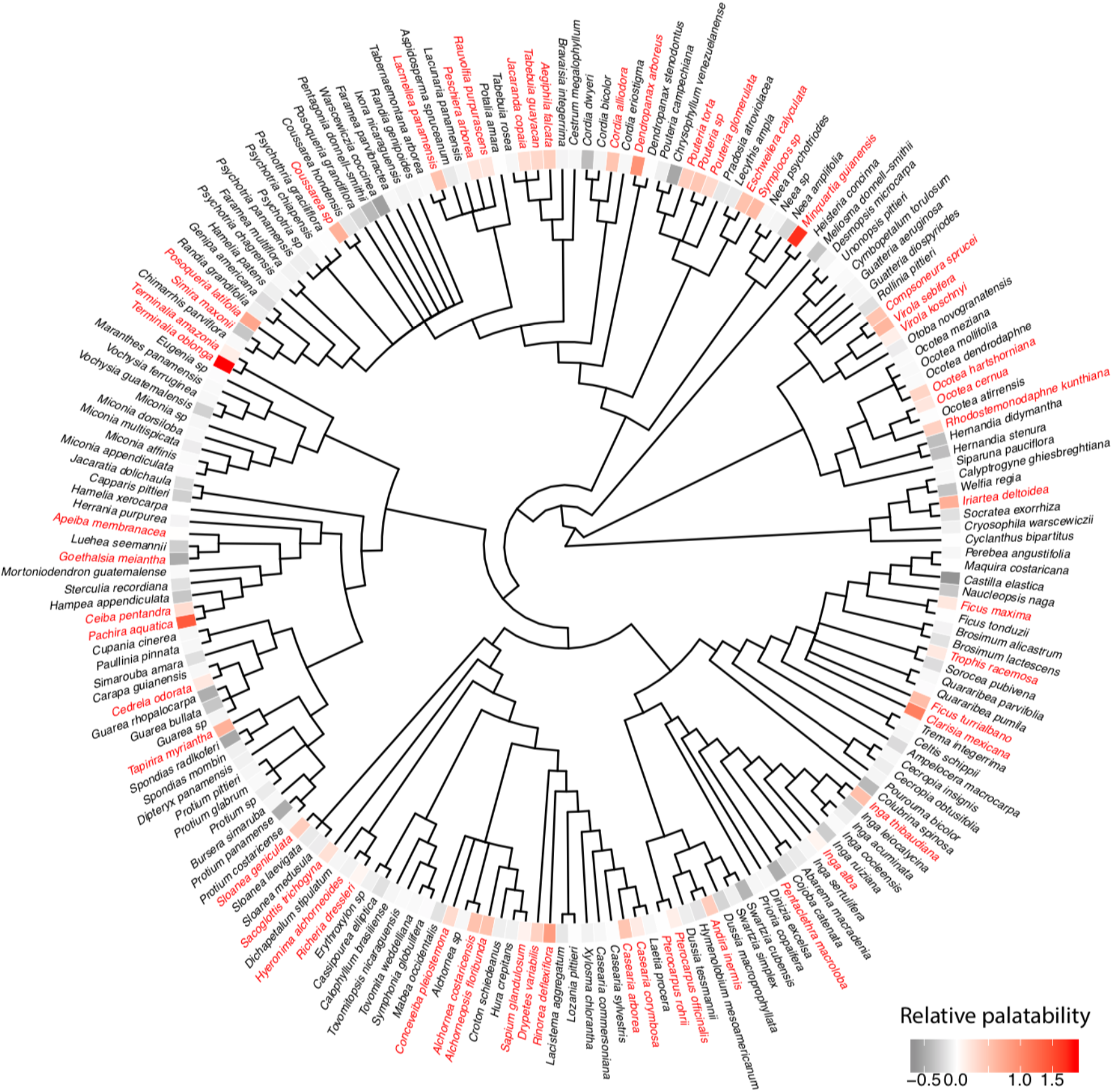
Phylogenetic tree of trees present in the arboretum (Figure 1), showing which species were foraged by leaf cutting ants. A statistically estimated degree of palatability for each species is given by a colored tile next to each species name. Red labels indicate species which were foraged at least once, while black indicates those which were not foraged in the course of this study. The model integrates repeated measures and phylogeny to estimate palatability coefficients, which are shown as a heatmap. Red colors indicate increasing palatability, while grey colours indicated lower palatability. Most species had zero or negative coefficients, suggesting that they were either not preferred or avoided by the ants. By contrast a relatively small number of trees species appeared to be actively preferred by the ants.

## Results

We indeed found evidence that all of them played a role in the likelihood of an individual tree being foraged (Table 1). However, it should be noted that the half of the variance remained unexplained (R^2^ = 0.50, 95% CI = 0.21-0.76). We used the same modelling approach without other covariates to estimate the strength of the phylogenetic signal, (equivalent to Pagel’s λ), which was estimated at 0.28 (95% CI = 0.13-0.39). In addition, taking advantage of phylogenetic interdependence between the tree species and the fact that several species were present more than one time, we could use these data to estimate the relative palatability of each species in the arboretum. The graphical summary of these analyses can be seen in Figure 2.

**Table 1.**
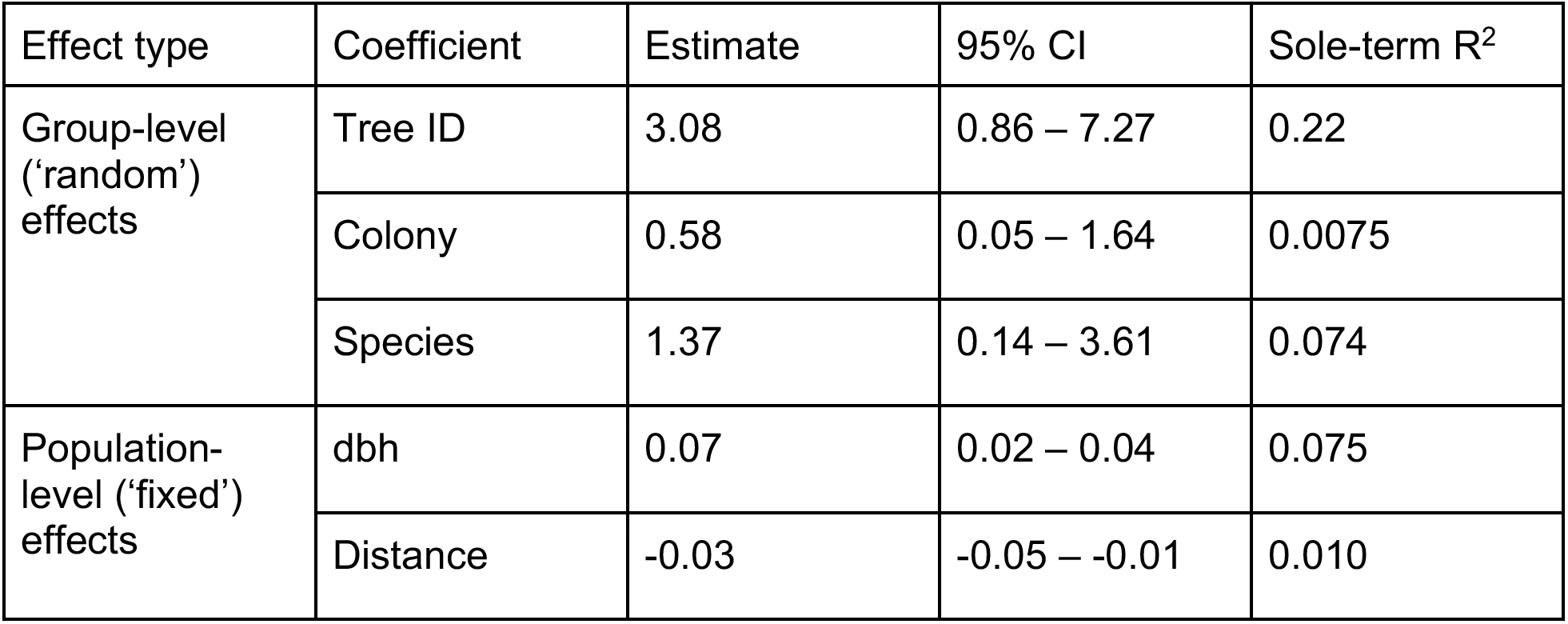
Effect sizes for the PGLS model. The table presents estimates and confidence intervals for the complete model as well as the R^2^ for the model with only the single term included as a way to compare their relative effect sizes. Foraging probability was positively affected by tree size, which could be due to greater biomass or a higher chance of finding palatable patches within a given tree. Trees farther from the nest mound were less likely to be foraged, possibly either because transport costs were higher, or because the trees at the edge of territories were contested between colonies (Figure 1). In addition, we found colony-level effects, and a phylogenetic effect on the likelihood of taking a particular tree species. Interestingly, the identity of the individual tree had the largest coefficient, suggesting that much of the variation in palatability came from tree-to-tree variability, rather than from species-level or colony-level effects, possibly due to variability present among individual trees in terms of chemical and physical characteristics resulting from factors such as age and life history. Interestingly, although most investigation has focused on ‘fixed’ effects, such as tree size and distance, ‘random’ effects explained a much greater percentage of the variance (R^2^ 0.096 *vs*. 0.525)

## Discussion

We found that the PGLS model captured essential features of LCA foraging behaviour, including the effects of tree size and distance to the colony (Costa *et al*. 2018, Farji-Brener *et al*. 2015, Kost *et al*. 2005). Yet, despite all the factors included in the model, there remained considerable unexplained variance. This suggests that there remain significant sources of variation to be explored. Some of them may be methodological and specific to the current study, such as the relatively short observation period, or the fact that the amount of forage collected was scored as a binary variable, which can lead to imprecise estimates. However, it is also likely that there may be other factors that were not included in the model that play major roles, for example, tree phenology, epiphyte cover of leaves, or other effects, which remain thus far undiscovered and should be included in future studies as explicit effects.

Up to now, most studies on leaf-cutting ant foraging have focused on testing specific well-defined hypotheses about how host plant properties affect ant behaviours, These have included tree size, the distance from colony to the foraging patch, as well as different measures of leaf characteristics (Farji-Brener *et al*. 2015, Rocha *et al*. 2017). Most often relatively easy-to-cut and least-defended leaves are preferred (Blanton & Ewel 1985, Coley & Barone 1996, Farji-Brener 2001, Wirth *et al*. 2003). Correspondingly, much research has focused on pioneer plant species (Fowler 1983, Shepherd, 1985, Farji Brener 2001, Wirth *et al*., 2003) and young leaves (Silva *et al*. 2013) which tend to have fewer chemical defences (Coley, 1983). Other aspects of foraging such as energy costs of cutting and transport while fulfilling the nutrient requirement and minimizing toxic intake as well have also been considered (Mundim et al. 2009). However, until now, there has not been a unifying statistical framework that could simultaneously address the relative importance of these effects. Furthermore, in addition to ‘fixed’ effects typically considered by most studies, uncontrolled ‘random’ effects may also explain much of the variation, which can be interesting in and of itself, or guide further investigations.

In this study we incorporated both fixed and random effects into the model, considering tree size and distance from tree to the colony as fixed effects and the individual tree identity, species identity of trees and the colony-level effect as random effects. According to our results, the two fixed effects have a small but non-negligible impact on foraging choice while the random effect is greater on foraging choice (Table 1). Incorporating different effect types allows direct comparisons between their effect sizes and can lead to surprising results. For example, the identity of each individual tree had the largest effect on foraging choice. This is in accordance with similar, previous studies which show that the foraging preference of leaf cutting ants are concentrated to a limited subset of host plant species (Cherrett 1968, Rockwood 1976). The observation that intraspecific variability has been previously shown by (Howard 1990) and is associated with leaf quality and physical characteristics. These include leaf moisture content, leaf toughness, leaf waxiness, presence of secondary chemicals and nutrient content, which varies both intra- and inter-specifically (Barrer & Cherrett 1972, Bowers & Porter 1981, Cherrett 1972, Cherrett & Seaforth 1970, Waller 1982). While much of the previous research has focused on interspecific variation, our results suggest that intraspecific variation may play a larger role in the ants’ foraging behaviour and deserves further investigation. We suggest that the intraspecific characteristics listed above, as well as other measurable factors of interest, be included in future studies as fixed effects. This will allow more specific partitioning of the variance and an assessment of their relative contribution to LCA dietary choices.

Our results broadly parallel a recent study by Gerhold *et al*. (2019), who also applied a phylogenetic approach to the study of *A. cephalotes* foraging in Brazil and likewise found λ = 0.25. However, the methods are not strictly comparable. For both their estimates of λ and a subsequent phylogenetically controlled analysis of leaf mechanical resistance, they used methods that expect continuous trait values, but palatability may have been scored as a binary character (Freckleton *et al*. 2002, Revell 2012, Orme *et al*. 2013). We believe that the approach presented here is a more general solution to the problem of analyzing field data on herbivore foraging because it allows the inclusion of (a) arbitrary fixed and random effects, and (b) the possibility of using discrete response variables. Furthermore, by taking advantage of the fact that multiple trees of the same species were present at the site, together with phylogenetic data, we were able to estimate ‘palatability coefficients’ for each species.

By incorporating multiple factors into a single model, the mixed-model approach allows comparison of overall model fit across datasets via the R^2^ statistic (Table 1). It would be interesting to re-analyze other data sets using the same approach to find out whether the per cent of the variance explained this study is unusually low, or whether our ability to predict LCA foraging behaviour remains low generally. In particular, it would be interesting to explore studies that quantified the amount of material consumed from each tree and species (Wirth *et al*. 1997), using a continuous rather than a binary response variable. Of course, it is also possible to apply this approach to other polyphagous herbivores, predators and parasites, as well as mutualists, such as pollinators. PGLS-based approaches should help understand what factors affect ecological interactions between species and even how they could have been affected by the evolutionary history of the host/prey.

## Acknowledgements

We are grateful to L. Elizalde, J. A. Meyers, S. C. Reed, S.A. Boyle and K. Proveda for their help in collecting data. This study began as a component of the Tropical Ecology field course with the Organization for Tropical Studies (OTS 03-01). Thanks to E. A. Herre, M. A. Kirkpatrick and the Barro Colorado Island Bambi seminar group for their comments and suggestions on an earlier version of the model.

